# Socioeconomic resources in youth are linked to divergent patterns of network integration and segregation across the brain’s transmodal axis

**DOI:** 10.1101/2023.11.08.565517

**Authors:** Cleanthis Michael, Aman Taxali, Mike Angstadt, Omid Kardan, Alexander Weigard, M. Fiona Molloy, Katherine L. McCurry, Luke W. Hyde, Mary M. Heitzeg, Chandra Sripada

**Author notes:** Corresponding Authors: Cleanthis Michael, Department of Psychology, University of Michigan, 2232 East Hall, 530 Church Street, Ann Arbor, MI 48109, USA, Chandra Sripada, Department of Psychiatry, University of Michigan, 4250 Plymouth Road, Ann Arbor, MI 48109, USA.

## Abstract

Socioeconomic resources (SER) calibrate the developing brain to the current context, which can confer or attenuate risk for psychopathology across the lifespan. Recent multivariate work indicates that SER levels powerfully influence intrinsic functional connectivity patterns across the entire brain. Nevertheless, the neurobiological meaning of these widespread alterations remains poorly understood, despite its translational promise for early risk identification, targeted intervention, and policy reform. In the present study, we leverage the resources of graph theory to precisely characterize multivariate and univariate associations between household SER and the functional integration and segregation (i.e., participation coefficient, within-module degree) of brain regions across major cognitive, affective, and sensorimotor systems during the resting state in 5,821 youth (ages 9-10 years) from the Adolescent Brain Cognitive Development (ABCD) Study. First, we establish that decomposing the brain into profiles of integration and segregation captures more than half of the multivariate association between SER and functional connectivity with greater parsimony (100-fold reduction in number of features) and interpretability. Second, we show that the topological effects of SER are not uniform across the brain; rather, higher SER levels are related to greater integration of somatomotor and subcortical systems, but greater segregation of default mode, orbitofrontal, and cerebellar systems. Finally, we demonstrate that the effects of SER are spatially patterned along the unimodal-transmodal gradient of brain organization. These findings provide critical interpretive context for the established and widespread effects of SER on brain organization, indicating that SER levels differentially configure the intrinsic functional architecture of developing unimodal and transmodal systems. This study highlights both sensorimotor and higher-order networks that may serve as neural markers of environmental stress and opportunity, and which may guide efforts to scaffold healthy neurobehavioral development among disadvantaged communities of youth.

## Introduction

Socioeconomic resources (SER) powerfully influence concurrent and lifelong outcomes, especially during childhood and adolescence when environmental experiences have strong and cascading effects on health and functioning (1–4). For example, household SER levels in youth, typically measured through family income, parental education, and neighborhood resources, have been associated with disparities in educational and occupational attainment, cognitive and socioemotional functioning, and physical (e.g., cardiovascular disease, cancer) and mental health (e.g., anxiety, depression, suicide, criminality, substance use) (5–9). Elucidating the biological mechanisms through which SER levels instigate pathways of vulnerability and resilience can inform early risk identification, facilitate targeted intervention, and encourage reform of public policies implicated in socioeconomic and mental health inequities.

Technological and computational advancements in non-invasive neuroimaging methods have allowed researchers to demonstrate that SER levels may influence behavior through their impact on brain function and development (10,11). Concurrently, there is increased recognition that the brain constitutes a complex network of interconnected regions (12,13). Task-free “resting-state” functional magnetic resonance imaging (fMRI) uses coherence in spontaneous activity across brain regions to yield maps of functional connectivity patterns that reflect neural communication within and between large-scale brain networks critical for cognition and mental health (14,15).

Previous studies probing how SER levels influence resting-state functional connectivity have predominantly relied on individual, region-specific connections (e.g., amygdala-ventromedial prefrontal connectivity) (16). There is, however, convergent evidence demonstrating that socioemotional, cognitive, and psychiatric characteristics emerge from widespread profiles of tens of thousands of connections across the entire brain, rather than focal profiles involving connections between individual pairs of regions (17).

Our group has therefore recently conducted the first multivariate predictive modeling study interrogating brain-wide connectivity changes associated with household SER (18) in the Adolescent Brain Cognitive Development (ABCD) Study, the largest neuroimaging study of youth to date (19,20). We identified robust and generalizable associations between SER and resting-state functional connectivity, with connectivity changes explaining 9% of the variance in SER out-of-sample – a relatively large effect size in the social sciences (21). These connectivity changes were widespread across most pairs of brain networks (72 out of 110 network pairs). A key limitation of this work, however, is in terms of interpretation. While we observed complex and widespread connectivity alterations associated with SER, the neurobiological meaning of these alterations remains elusive.

In the present study, we address this knowledge gap by leveraging the resources of graph theory (22). The human brain is organized into multiple intrinsic connectivity networks (ICNs) (23–25). ICNs exhibit developmental refinements in profiles of segregation (i.e., the degree of neural communication within distinct, functionally specialized networks) and integration (i.e., the degree of neural communication across different networks) during sensitive developmental windows (26–29). Integration and segregation are reflected in a pair of graph theoretic metrics that describe between-network connectivity (participation coefficient) and within-network connectivity (within-module degree) (30). Profiles of higher participation coefficient and lower within-module degree reflect integration, while the reverse reflects segregation (31).

ICNs are organized along a unimodal-transmodal gradient, which represents the degree to which networks are specialized for encoding specific sensory features versus integrating representations across modalities (32–34). Motor and sensory processing networks anchor the unimodal end, heteromodal networks occupy the middle range, and association networks anchor the transmodal end (32–34). Across development, unimodal networks become more integrated and transmodal networks become more segregated (27,29). SER levels have been previously associated with functional network integration/segregation in youth (35–37). As different ICNs exhibit unique developmental refinements based on their position on the sensorimotor-association gradient, the topological effects of SER may differ along the transmodal axis, though this possibility currently remains unclear.

Multiple ecological mechanisms associated with SER (e.g., parental stimulation, school quality, nutrition, neighborhood adversity) may influence coordinated patterns of ICN organization, especially in terms of integration and segregation (11,38). Thus, in the present study, we quantify multivariate and univariate associations between household SER and the within-module degree and participation coefficient of 418 nodes across 15 major ICNs throughout the brain. Moreover, we assess potential ICN-specific effects of SER (e.g., greater segregation and lower integration in certain networks; the reverse in others). Finally, we interrogate whether the effects of SER on network integration/segregation are spatially patterned along the sensorimotor-association axis.

We performed our analyses in the ABCD Study, a population-based consortium study of 11,875 9- and 10-year-olds with substantial sociodemographic diversity (39). As in our prior report (18), we constructed a latent factor of SER across household and neighborhood contexts. We establish that SER has robust relationships with network integration/segregation, which account for most of the association between SER and the entire functional connectome. Furthermore, we delineate network-specific effects, with higher SER related to greater integration of sensorimotor networks but greater segregation of association networks. Lastly, we demonstrate that the effects of SER strongly relate to the transmodal axis. These findings add valuable interpretive information by suggesting that the associations between SER and functional connectivity spatially conform to the sensorimotor-association axis during development. Such insights may elucidate neural markers of environmental stress and opportunity. Moreover, they may guide interventions that support patterns of brain organization linked to enhanced executive functioning and emotional wellbeing during early adolescence, a critical window when many psychosocial challenges emerge (26,28,40).

## Materials and Methods

### 1. Sample and Data

The ABCD Study is a multisite longitudinal study with 11,875 children between 9-10 years of age from 22 sites across the United States. The study conforms to the rules and procedures of each site’s Institutional Review Board, and all participants provide informed consent (parents) or assent (children). Data for this study are from ABCD Release 3.0.

### 2. Data Acquisition, fMRI Preprocessing, and Connectome Generation

High spatial (2.4mm isotropic) and temporal resolution (TR = 800ms) resting-state fMRI was acquired in four separate runs (5min per run, 20min total). Preprocessing was performed using fMRIPrep v1.5.0 (41). Briefly, T1-weighted (T1w) and T2-weighted images were run through recon-all using FreeSurfer v6.0.1, spatially normalized, rigidly coregistered to the T1, motion corrected, normalized to standard space, and transformed to CIFTI space.

Connectomes were generated for each functional run using the Gordon 333 parcel atlas (42), augmented with parcels from high-resolution subcortical (43) and cerebellar (44) atlases. Volumes exceeding a framewise displacement (FD) threshold of 0.5mm were marked to be censored. Covariates were regressed out of the time series in a single step, including: linear trend, 24 motion parameters (original translations/rotations + derivatives + quadratics), aCompCorr 5 CSF and 5 WM components and ICA-AROMA aggressive components, high-pass filtering at 0.008Hz, and censored volumes. Next, correlation matrices were calculated. Full details of preprocessing and connectome generation are reported in the Supplement and the automatically-generated fMRIPrep Supplement.

### 3. Inclusion/Exclusion

There are 11,875 subjects in the ABCD Release 3.0 dataset. Subjects were excluded for: failing ABCD QC, insufficient number of runs each 4 minutes or greater, failing visual QC of registrations and normalizations, and missing data required for regression modeling. This left us with *N* = 5,821 subjects across 19 sites for the main analysis. Details of exclusions are provided in the Supplement.

### 4. Graph Theoretic Analysis

Since most graph theory measures require unsigned edge weights, each subject’s connectome resulted in two separate sets of graphs – one for the collection of positive edges and another for the negatively weighted edges (45,46). We focused on positive graphs consistent with previous graph theoretical investigations (45,46), though supplementary analyses revealed that negative graphs did not add meaningful predictive information (see Supplement).

*Within-Module Degree* is a node-wise measure which captures each node’s degree (i.e., the magnitude of summed connectivity weights) specifically within the node’s own network. This measure is a modification of the “module degree z-score” metric (30), but without within-network z-scoring of node degree to better capture differences across participants, rather than differences across nodes within each network. Formally, the within-module degree of a node *i* is given by:

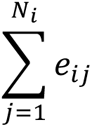

where *e_ij_* is the edge weight between nodes *i* and *j*, and *N_i_* is the set of nodes incident to node *i* that are in the same network as *i*.

Participation Coefficient is a node-wise measure that captures the diversity of a node’s connections with other nodes outside of its own network (30). Intuitively, if a node distributes its connectivity evenly across all networks, its participation coefficient will be 1, while departures from equality yield commensurately lower scores. Formally, the participation coefficient of a node i is given by:

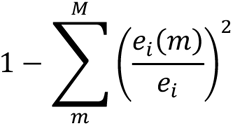

where *M* is the set of networks, *e_i_*(*m*) is the sum of edge weights between node *i* and all nodes in network *m* and *e_i_* is the sum of edge weights between node *i* and all other nodes.

For both metrics, we used the community structure defined by the applied parcellation schemes to determine network boundaries. Within-module degree (MDP) and participation coefficient (PCP) for positive edges were calculated for 418 nodes, yielding 836 node-wise graph theoretic features per participant.

To quantify the multivariate relationship between these 836 graph theoretic metrics and SER, we used principal component regression (PCR) predictive modeling (47,48) (Figure S2). Briefly, this method performs dimensionality reduction on the set of predictive features (i.e., graph theoretic metrics), fits a regression model on the resulting components (where the number of components is determined in nested cross-validation), and applies this model out-of-sample in a leave-one-site-out cross-validation framework. We control for multiple nuisance covariates, including sex assigned at birth, parent-reported race/ethnicity, age, age-squared, mean FD, and mean FD-squared. We controlled for race/ethnicity, a social construct, to account for differences in exposure to personal/systemic racism, disadvantage, and opportunity among people of color (49,50). We assessed statistical significance with non-parametric permutation tests, using the procedure of Freedman and Lane (51) to account for covariates. Exchangeability blocks were used to account for twin, family, and site structure and were entered into Permutation Analysis of Linear Models (PALM) (52) to produce permutation orderings. Details on these analyses are provided in the Supplement.

### 5. Latent Variable Modeling

We constructed a latent variable for SER by applying exploratory factor analysis to household income-to-needs, parental education, and neighborhood disadvantage (18). Household income-to-needs represents the ratio of a household’s income relative to its need based on family size (details provided in the Supplement). Parental education was the average educational achievement of parents or caregivers. Neighborhood disadvantage scores reflect an ABCD consortium-supplied variable (reshist_addr1_adi_wsum). In brief, participants’ primary home address was used to generate Area Deprivation Index (ADI) values (53), which were weighted based on results from Kind et al. (54) to create an aggregate measure. Additional details on construction of this latent variable are provided in the Supplement.

### 6. Code Availability

The ABCD data used in this report came from NDA Study 901, 10.15154/1520591, which can be found at https://nda.nih.gov/study.html?id=901. The subsample used for this study can be found at NDA DOI: 10.15154/ebhq-f780. Code for running analyses can be found at https://github.com/SripadaLab/ABCD_Resting_SER_GraphTheory.

## Results

### 1. Within-module degree and participation coefficient are strongly related to household socioeconomic resources

As reported in our previous study (18), using leave-one-site-out cross-validation (LOSO-CV), the out-of-sample multivariate relationship between SER and the whole connectome (reflecting 87,153 connections) was r_cv_ = 0.274, p_PERM_ < 0.0001. Against this benchmark result, we found that the LOSO-CV out-of-sample multivariate relationship between SER and these 836 node-wise graph theoretic measures (i.e., MDP, PCP) was r_cv_ = 0.162, p_PERM_ < 0.0001. Thus, the linear MDP/PCP-SER relationship is 59.1% as strong as the whole connectome-SER relationship.

We next examined whether the 836 MDP/PCP features reflect distinct or overlapping variance in predicting SER relative to the 87,153 connections of the entire functional connectome. To assess this, we built a stacked model by taking the SER predictions from the full connectome predictive model, and the MDP/PCP predictive model, and entering them as predictors of SER in a new regression. This stacked model’s LOSO-CV out-of-sample performance was r_cv_= 0.268; that is, the stacked model with the addition of graph theory features performed no better than the full connectome model by itself.

These results suggest two conclusions. First, the graph theoretic features represent a subset of the variance explained by the whole connectome. Second, there is strong concentration of SER predictivity in the graph theoretic features, wherein these 836 graph theoretic features account for the majority of the multivariate relationship between the functional connectome and SER.

### 2. Associations between household socioeconomic resources and patterns of integration/segregation differ across intrinsic connectivity networks

In Figure 1, we display standardized regression weights for 418 node-wise MDP metrics and 418 PCP metrics, each regression weight arising from separate regression models predicting SER from the respective metric (with controls for nuisance covariates). The plot highlights strongly divergent relationships with SER across different ICNs, with four notable zones. Zone 0 contains the majority of nodes that lack statistically significant relations with SER. In Zone 1, we observe large, positive SER-predictive betas for MDP in default mode network, an unlabeled network (dubbed “None”) primarily anchored in orbitofrontal cortex, and cerebellum, indicating greater segregation of nodes within these networks with higher SER. In Zone 2, we observe large, positive SER-predictive betas for PCP primarily in subcortical networks, indicating greater integration of nodes within this network with higher SER. In Zone 3, we observe large SER-predictive betas for both MDP (negative betas) and PCP (positive betas) primarily with nodes in the somatomotor network, indicating greater integration of nodes within this network with higher SER.

**Figure 1:**
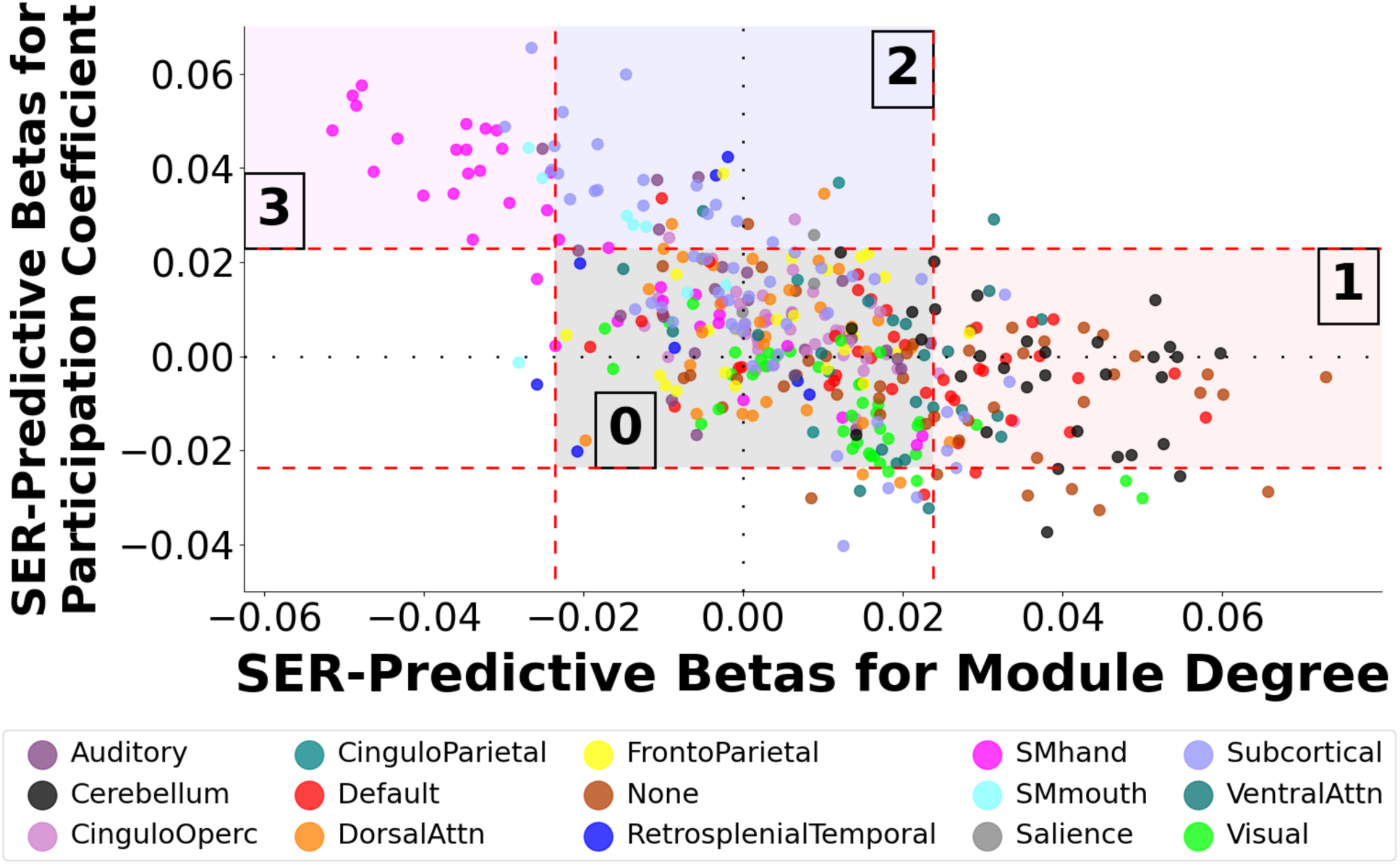
Profile Plot Showing Relation Between Within-Module Degree and Participation Coefficient Beta Weights When Predicting Socioeconomic Resources by Network Affiliation. We computed beta weights from 836 regression models in which socioeconomic resource (SER) scores were the outcome variable predicted by 418 metrics of within-module degree for positive edges (MDP) and 418 metrics of participation coefficient for positive edges (PCP). Each node’s pair of SER-predictive betas (for MDP and PCP) is shown in the above “profile plot”, with nodes shaded by network affiliation. Orange lines represent the thresholds for statistically significant univariate relationships between SER and MDP/PCP metrics. Four zones are noteworthy. Zone 0 contains the majority of nodes that lack statistically significant relations with SER. Zone 1 nodes exhibit positive SER-predictive betas for MDP, consistent with greater segregation of these nodes with higher SER. Zones 2 and 3 exhibit higher SER-predictive betas for PCP (Zone 2 and 3) and lower SER-predictive betas for MDP (Zone 3), consistent with greater integration of these nodes with higher SER. Somatomotor-hand, in the upper left, stands out as exhibiting particularly extensive integration with higher SER. CinguloOperc = Cingulo-Opercular Network. DorsalAttn = Dorsal Attention Network. SMhand = Somatomotor Hand Network. SMmouth = Somatomotor Mouth Network. VentralAttn = Ventral Attention Network.

**Figure 2:**
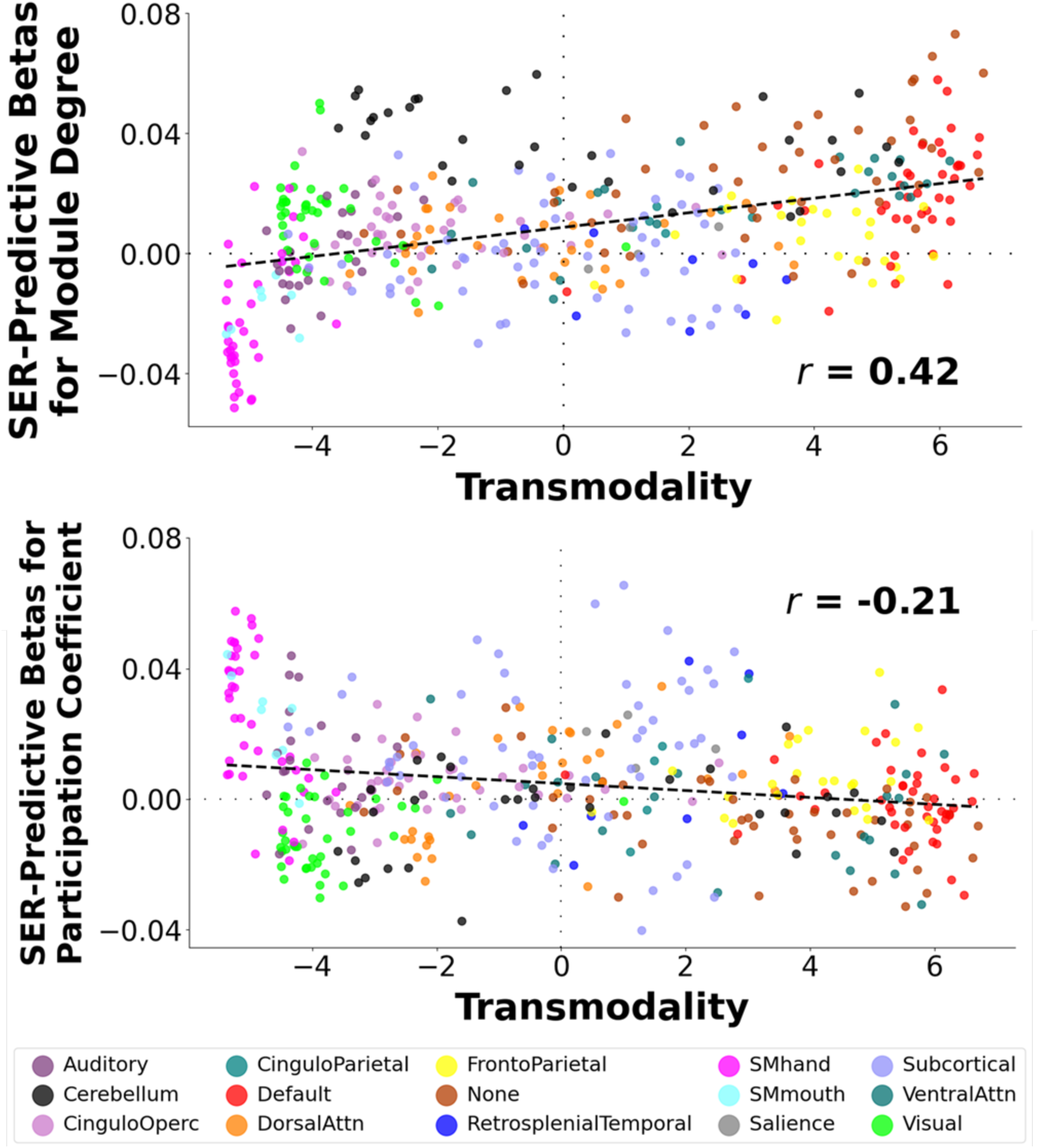
Scatter Plots Showing Relationships between Within-Module Degree and Participation Coefficient Beta Weights When Predicting Socioeconomic Resources with Transmodality Scores. Top Figure: We obtained transmodality scores from 418 nodes from a previous report by Margulies and colleagues (34), which locates nodes along a gradient with sensory processing networks at one end (lowest transmodality scores) and higher-order association networks at the other end (highest transmodality scores). In addition, we calculated associations between within-module degree for positive edges (MDP) scores for each of these nodes and socioeconomic resources (SER) (“SER-predictive betas for MDP”). We found a strong positive association between transmodality scores and SER-predictive betas for MDP. Bottom Figure: We performed this same analysis, but this time with participation coefficient for positive edges (PCP) scores. We found a moderate negative association between transmodality scores and SER-predictive betas for PCP. Nodes shaded by network affiliation. CinguloOperc = Cingulo-Opercular Network. DorsalAttn = Dorsal Attention Network. SMhand = Somatomotor Hand Network. SMmouth = Somatomotor Mouth Network. VentralAttn = Ventral Attention Network.

### 3. Household socioeconomic resource levels exhibit divergent relationships with network integration/segregation across the brain’s unimodal-transmodal gradient

Given differences in the segregation/integration profiles of different ICNs in relation to SER, we next examined whether these differences are associated with the transmodality axis. We used transmodality scores from a previous report by Margulies and colleagues (34), which locates nodes along a gradient with sensory processing networks at one end (lowest transmodality scores) and higher-order association networks at the other end (highest transmodality scores). We found that transmodality scores exhibited a strong positive relationship with SER-predictive betas for MDP (r = 0.42, p_PERM_ < 0.007), and a moderate negative relationship with SER-predictive betas for PCP (r = 0.21) that only trended toward significance (p_PERM_ < 0.09). These results provide quantitative support for divergent SER effects across the transmodality gradient, with SER yielding greater integration (lower MDP regression weights, higher PCP regression weights) at the sensorimotor processing pole and greater segregation (higher MDP regression weights, lower PCP regression weights) at the higher-order processing pole.

## Discussion

Household SER levels across childhood and adolescence calibrate structural and functional neurodevelopment, with potent implications for physical health, occupational attainment, and emotional wellbeing across the lifespan (10,11,55). In the present report, we leverage graph theory and the largest neuroimaging cohort of youth to date to delineate how variation in household SER becomes biologically expressed along the developing functional architecture of cognitive, affective, and sensorimotor brain systems. We found that SER was robustly associated with two graph theoretic metrics that decompose brain organization in terms of integration and segregation. Importantly, the topological effects of SER were not uniform across the brain; rather, higher SER levels were related to greater integration of somatomotor and subcortical systems, but greater segregation of default mode, orbitofrontal, and cerebellar systems. Finally, we demonstrate that SER-related network reconfiguration was spatially patterned along the brain’s transmodal axis. These findings provide critical interpretive context for the established and widespread effects of SER on the intrinsic functional architecture of the developing brain.

Previous studies characterizing the neurobiological embedding of SER have primarily examined connections between individual pairs of regions (e.g., frontolimbic connectivity) (16). Given the brain-wide effects of SER (36,37,56), and the thousands of connections that undergird complex and clinically relevant phenotypes (17), our group recently conducted the first multivariate predictive modeling study of household SER in the ABCD Study (18). We revealed that the correlation between actual SER and SER predicted from 87,153 functional connections at rest was 0.27, yet the neuroscientific meaning of these findings remained unclear. In this study, we applied graph theory to distill these 87,153 connections into only 836 features that describe the effects of SER with greater neurobiological interpretability in terms of intra- and inter-network relationships. Specifically, we assessed node-level integration and segregation using participation coefficient (between-network connectivity) and within-module degree (within-network connectivity), and we demonstrate that these two metrics capture more than half of the original association with SER (r = 0.16). These findings indicate that these two nodal graph properties largely capture the backbone of functional brain architecture, particularly in relation to SER.

Segregation gives rise to differentiated networks that execute specialized cognitive functions, whereas integration efficiently coordinates these processing streams across the brain (57,58). A combination of high segregation and high integration represents an “optimized” small-world architecture that rapidly integrates specialized, multimodal information at low wiring and energy costs (59,60). Our multivariate findings therefore suggest that the developmental construction of an “optimal” small-world-like configuration may be impacted by SER.

To spatially localize the topological effects of SER, we next conducted univariate analyses probing the within-module degree and participation coefficient of brain regions within 15 major ICNs. First, we found that higher SER levels were associated with greater segregation (higher within-module degree) of the default mode network, an unlabeled network (dubbed “None”) primarily anchored in orbitofrontal cortex, and the cerebellum. These systems are respectively purported to support self-referential and introspective cognition, reward processing and decision-making, and cognitive and motor control (25,61–63) and have been previously linked with SER, despite some inconsistencies in directionality (36,37,56,64,65). As segregation of these systems is associated with attention, cognitive control, and impulsivity (62,66,67), these alterations may represent a mechanistic pathway from socioeconomic gradients to goal-directed, regulatory behavior in youth.

Second, we found that higher SER scores were associated with greater functional integration (higher participation coefficient) of the subcortical network implicated in motor planning, threat and safety learning, and emotion processing (68–71). These findings converge with extensive evidence linking SER to structural, functional, and connectivity profiles of subcortical regions, such as the amygdala and hippocampus (72–74). Given their dense expression of glucocorticoid receptors (75,76), these structures may be particularly sensitive to both nurturing and stressful experiences often associated with SER (11,77). Integration of subcortical regions with cortical systems subserves adaptive emotional learning and regulation (71,78), indicating a plausible network-level neural basis for documented links between poverty and psychopathology (3,6,8).

Lastly, higher SER levels were strongly associated with greater functional integration (lower within-module degree, higher participation coefficient) of the somatomotor hand network. This network is not commonly considered in theoretical accounts linking SER to brain development (10,11,79,80), despite being consistently implicated in SER and transdiagnostic psychopathology in individual studies (36,56,81,82). The somatomotor network supports motor planning and execution (25), and recent data point to its potential involvement in a “somato-cognitive action” network that integrates motoric function with goal-directed planning (83). One possibility is that SER levels not only calibrate association systems that generate and evaluate abstract cognitive representations, but also somatomotor systems that translate these abstract representations into goal-relevant behavior. These findings highlight the need for theoretical accounts and empirical studies to further delineate how adversity constrains or reconfigures somatomotor development to confer vulnerability and resilience.

Since SER displayed divergent associations with the integration/segregation of different ICNs, we investigated whether this heterogeneity could be explained by considering how ICNs are organized along the brain’s unimodal-transmodal axis. This evolutionarily rooted, hierarchical axis of brain organization is anchored by sensory and motor networks on one end and association networks on the other (32–34). This sensorimotor-association gradient captures developmental sequences of multiple neurobiological properties, from structure and myelination to plasticity and gene expression (26,84). In the present investigation, we hypothesized that this axis may also provide a unifying framework for characterizing the network-specific effects of household SER. Consistent with this hypothesis, we found that associations between SER and functional network integration/segregation were indeed spatially patterned along the transmodal axis, with higher SER levels associated with greater integration at the unimodal/somatosensory pole and greater segregation at the transmodal/association pole.

Over the course of neurodevelopment from childhood to young adulthood, lower-order unimodal networks (e.g., somatomotor network) become more integrated, whereas higher-order association networks (e.g., default mode network) become more segregated (27,29). Thus, the construction of integrated somatomotor systems and segregated association systems may represent a universal milestone of functional neurodevelopment. Against this backdrop, our findings suggest that higher SER may facilitate the emergence of this sensorimotor-association hierarchy. Conversely, lower SER may predict developmental lags in the emergence of this configuration, consistent with cross-sectional and longitudinal findings suggesting disadvantage-related delays in the pace of neurodevelopment (37,74,85–88). Candidate mechanisms for protracted brain development following disadvantage include material hardship (e.g., resource access, lower-quality nutrition), less complex social and cognitive stimulation (e.g., under-resourced schools, complex reading materials), and toxicant exposure (e.g., lead, particulate matter) (11,86). These exposures may alter synaptic proliferation and pruning, and ultimately maturational refinements in functional network communication (integration) and specialization (segregation) (89–91),

Nevertheless, an alternative interpretation of our findings is that developmental trajectories and milestones of brain organization may differ as a function of household SER. In other words, the trajectory and outcome of neurodevelopment may be qualitatively different depending on SER. While higher-SER youth may establish an integrated unimodal and segregated transmodal pole with development, lower-SER youth may develop distinct profiles of integration/segregation. These distinct neural profiles may allow youth to successfully navigate the unique demands of disadvantaged environments but may also manifest in cognitive and socioemotional challenges across the lifespan. The former hypothesis is consistent with data indicating that functional connectivity patterns that optimize cognition differ in high-versus low-SER contexts (92), as well as a recent review of longitudinal studies concluding that disadvantage may engender unique, rather than temporally shifted, trajectories of structural brain development (91).

In a separate report (in preparation), we repeated our analyses evaluating associations between sleep duration, rather than SER, with the functional integration/segregation of the same 15 ICNs in the ABCD Study. Strikingly, we found that sleep duration displayed similar but even stronger associations with functional network architecture. Consistent with the reported effects of SER, these associations were strongest for the organization of the somatomotor network, such that youth who sleep for a longer duration exhibit a more integrated somatomotor network. These findings accord with recent studies linking sleep quality to somatomotor connectivity (93–95) and suggest that somatomotor architecture may represent a robust neural marker associated with multiple forms of environmental stress, adversity, and opportunity during development.

Our study has several limitations that will be important to address in future research. First, our analyses are cross-sectional and thus do not support inferences about the direction of causality of associations or about patterns of neurodevelopment. As neuroimaging data from future ABCD waves are released, future studies should disentangle causal effects and assess how the spatially divergent effects of SER unfold longitudinally across development. Second, SER scores in the ABCD Study are overall higher compared to the national population, an issue that is further exacerbated by our exclusion criteria (e.g., cutoffs for excessive head motion) (96,97); thus, caution should be exercised when attempting to generalize our findings to the broader population in the United States and worldwide. Lastly, in our previous multivariate study of SER (18), granular analyses demarcated that parental education was the primary factor related to functional connectivity (compared to family income-to-needs and neighborhood disadvantage). Here, our focus is on interpreting and spatially localizing these multivariate effects. This focus introduces challenges in dissecting the unique role of each SER component, which constitutes an important future direction to inform priorities for policy, prevention, and intervention.

In sum, the present study provides essential neuroscientific meaning to the established and widespread effects of household SER on intrinsic functional connectivity. By integrating methodological advancements in network neuroscience with theoretical frameworks of brain organization, we demonstrate that associations between SER and profiles of network integration/segregation in youth unfold differentially along the brain’s transmodal axis, with stronger effects on default mode, cerebellar, subcortical, and somatomotor networks. Our findings illustrate that SER levels may calibrate the intrinsic graphical architecture of the developing brain, highlighting the importance of prevention and intervention efforts that facilitate the development of cognitive, affective, and sensorimotor processes underlying risk and resilience within disadvantaged communities of youth.

## Acknowledgements

Data used in the preparation of this article were obtained from the Adolescent Brain Cognitive Development (ABCD) Study (https://abcdstudy.org), held in the NIMH Data Archive (NDA). This is a multisite, longitudinal study designed to recruit more than 10,000 children aged 9-10 years and follow them over 10 years into early adulthood. The ABCD Study is supported by the National Institutes of Health and additional federal partners under award numbers U01DA041022, U01DA041028, U01DA041048, U01DA041089, U01DA041106, U01DA041117, U01DA041120, U01DA041134, U01DA041148, U01DA041156, U01DA041174, U24DA041123, and U24DA041147. A full list of supporters is available at https://abcdstudy.org/nih-collaborators. A listing of participating sites and a complete listing of the study investigators can be found at https://abcdstudy.org/principal-investigators.html. ABCD consortium investigators designed and implemented the study and/or provided data but did not necessarily participate in analysis or writing of this report. This manuscript reflects the views of the authors and may not reflect the opinions or views of the NIH or ABCD consortium investigators. The ABCD data repository grows and changes over time. The ABCD data used in this report came from NDA Study 901, 10.15154/1520591, which can be found at https://nda.nih.gov/study.html?id=901.

## Funding

AW was supported by K23 DA051561 and R21 MH130939. KLM, OK, and MFM were supported by T32 AA007477. CS was supported by R01 MH123458 and U01DA041106.

## Conflict of Interest

The authors declare that they have no known competing financial interests or personal relationships that could have appeared to influence the work reported in this paper.

## Notes

### Competing Interest Statement

The authors have declared no competing interest.

