## Supplementary Material for "Socioeconomic resources in youth are linked to divergent patterns of network integration and segregation across the brain’s transmodal axis"

### Supplemental Methods and Results

#### 1. Sample and Data

The ABCD study is a multisite longitudinal study with 11,875 children between 9-10 years of age from 22 sites across the United States. The study conforms to the rules and procedures of each site's Institutional Review Board, and all participants provide informed consent (parents) or assent (children). Data for this study are from ABCD Release 3.0.

#### 2. Data Acquisition, fMRI Preprocessing, and Connectome Generation

Imaging protocols were harmonized across sites and scanners. High spatial (2.4mm isotropic) and temporal resolution (TR = 800ms) resting-state fMRI was acquired in four separate runs (5min per run, 20min total). The entire data pipeline described below was run through automated scripts on the University of Michigan's high-performance cluster, and is described below.

Preprocessing was performed using fMRIPrep version 1.5.0 (1), and detailed methods automatically generated by fMRIPrep software are provided in the fMRIPrep Supplement. T1-weighted (T1w) and T2-weighted images were run through recon-all using FreeSurfer v6.0.1. T1w images were also spatially normalized nonlinearly to MNI152NLin6Asym space using ANTs 2.2.0. Each functional run was corrected for fieldmap distortions, rigidly coregistered to the T1, motion corrected, and normalized to standard space. ICA-AROMA was run to generate aggressive noise regressors. Anatomical CompCor was run and the top 5 principal components of both CSF and white matter were retained. Functional data were transformed to CIFTI space using HCP's Connectome Workbench. All preprocessed data were visually inspected at two separate stages to ensure only high-quality data was included: after co-registration of the functional data to the structural data and after registration of the functional data to MNI template space.

Connectomes were generated for each functional run using the Gordon 333 parcel atlas (2), augmented with parcels from high-resolution subcortical (3) and cerebellar (4) atlases. Volumes exceeding a framewise displacement threshold of 0.5mm were marked to be censored. Covariates were regressed out of the time series in a single step, including: linear trend, 24 motion parameters (original translations/rotations + derivatives + quadratics), aCompCorr 5 CSF and 5 WM components and ICA-AROMA aggressive components, high-pass filtering at 0.008Hz, and censored volumes. Next, correlation matrices were calculated for each run. Each matrix was then Fisher r-to-z transformed, and then averaged across runs for each subject yielding their final connectome.

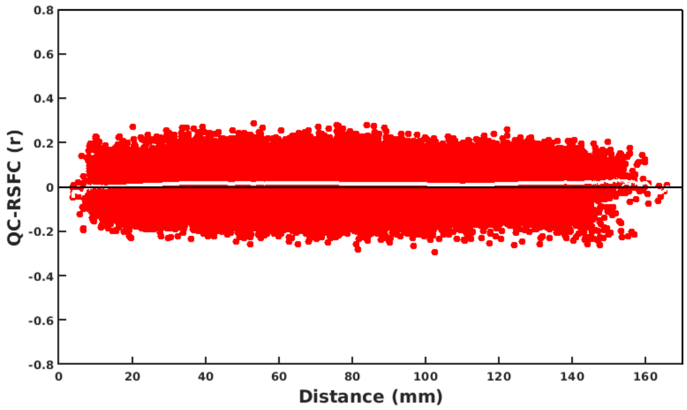


***Figure S1: Quality Control-Resting State Functional Connectivity Plot***

We used multiple procedures listed above to limit the effect of head motion on resting-state functional connectivity maps. To assess the effectiveness of these procedures, we produced a quality control resting-state functional connectivity (QC-RSFC) plot (5,6). This plot shows the relationship between mean framewise displacement and connectivity edges binned by distance. Motion effects produce a sloped line (distance-dependent artifact), while a flat line is indicative of minimal motion-related effects. The QC-RSFC plot for our ABCD resting-state data showed a flat line (Figure S1), providing additional evidence that our stringent motion correction strategies were effective.

#### 3. Inclusion/Exclusion

There are 11,875 subjects in the ABCD Release 3.0 dataset. Screening was initially done using ABCD raw QC to limit to subjects with 2 or more good runs of resting data as well as a good T1 and T2 image (QC score, protocol compliance score, and complete all = 1). This resulted in 9,580 subjects with 2 or more runs that entered preprocessing. Each run was subsequently visually inspected for registration and warping quality, and only those subjects who still had 2 or more good runs were retained (N = 8,858). After connectome generation, runs were excluded if they had less than 4 minutes of uncensored data, and next subjects were retained only if they had 2 or more good runs (N = 6,568). Finally, subjects who were missing data required for factor modeling of socioeconomic resources (SER) were dropped and sites with fewer than 75 subjects were dropped. This left us with N = 5,821 subjects across 19 sites for the neuroimaging analysis, and demographic characteristics of the overall and included sample are shown in Table S1.

| | Included | Excluded |
|---|---|---|
| N | 5821 | 6057 |
| Age (mean (s.d.)) | 9.96 (0.62) | 9.87 (0.62) |
| Female (%) | 2920 (50.2) | 2762 (45.6) |

| | | |
|---|---|---|
| Race-Ethnicity (%) | | |
| non-Hispanic White | 3469 (59.6) | 2710 (44.7) |
| non-Hispanic Black | 667 (11.5) | 1115 (18.4) |
| Hispanic | 1052 (18.1) | 1359 (22.4) |
| non-Hispanic Asian | 86 (1.5) | 187 (3.1) |
| Multi-racial/Other | 547 (9.4) | 686 (11.3) |
| No answer | -- | -- |
| Highest Parental Education (%) | | |
| < HS Diploma | 177 (3.0) | 416 (6.9) |
| HS Diploma/GED | 406 (7.0) | 726 (12.0) |
| Some College | 1461 (25.1) | 1619 (26.7) |
| Bachelor | 1614 (27.7) | 1401 (23.1) |
| Post Graduate Degree | 2163 (37.2) | 1881 (31.1) |
| No answer | 0 (0.0) | 14 (0.2) |
| Household Marital Status – Married (%) | 4202 (72.2) | 3789 (62.6) |
| Household Income (%) | | |
| <50K | 1585 (27.2) | 1944 (32.1) |
| >=50k & <100K | 1723 (29.6) | 1387 (22.9) |
| >=100k | 2513 (43.2) | 2082 (34.4) |
| No answer | 0 (0.0) | 644 (10.6) |

***Table S1: Demographic Characteristics of Included Versus Excluded Subjects***

#### 4. Principal Components Regression-Based Multivariate Predictive Modeling

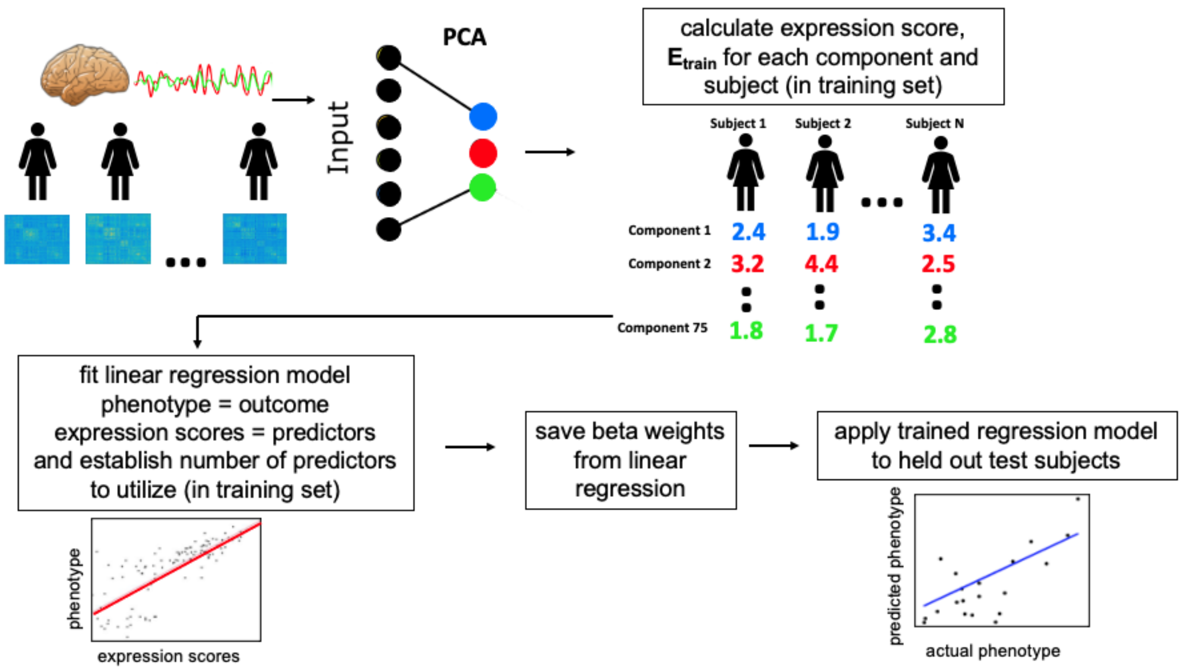


***Figure S2: Steps of Principal Component Regression Predictive Modeling.***

We implemented principal component regression (PCR) (7) as a multivariate predictive modeling method for identifying brain-behavior relationships (8) (see Figure S2). The method involves two key steps: 1) Use principal component analysis (PCA) to find a set of components that capture *inter-individual* differences in brain features; 2) Use multiple regression in a cross-validation framework to link expression scores for these components to phenotypes of interest. In previous work, we often used the more general name brain basis set (BBS) for this approach to capture commonalities with work by our group and others that use alternative methods for step 1 (e.g., independent component analysis (9,10) or community detection (11,12)). We chose the PCR approach for this study because our previous work showed it has high test-retest reliability (13) and predictive accuracy (14,15) and generally performs as well as or better than alternative methods such as support vector regression and ridge regression (13).

We performed PCA dimensionality reduction on an *n* subjects by *p* connectivity features matrix, yielding *n* principal components (i.e., directions in the feature space) that represent inter-individual differences in the imaging features (functional connectivity or graph theory metrics). Pre-subject expression scores for a subset of *k* of these components then entered multiple regression modeling to identify linear associations with phenotypes of interest (here, SER scores). Of note, we selected *k* using 5-fold cross-validation within the training data, as in our previous work (13).

To assess accuracy and generalizability of PCR predictive models, we used leave-one-site-out cross-validation. In each fold of the cross-validation, data from one of the 19 sites served as the held-out test dataset and data from the other 18 sites served as the training dataset. Additionally, to ensure separation of train and test datasets, at each fold of the cross-validation, a new PCA was performed on the imaging features (functional connectivity or graph theory metrics) in the training dataset, and expression scores of these brain components were calculated for the test set. Note that by employing leave-one-site-out, members of twinships and sibships are never present in both training and test samples. We assessed the performance of PCR predictive models with cross-validated Pearson's correlation and cross-validated partial eta squared.

In each fold of the leave-one-site out cross-validation (LOSO-CV), PCR predictive models were trained in the train partition with the following covariates (unless explicitly stated otherwise for specific analyses): sex, race-ethnicity, age, age squared, mean FD and mean FD squared. To maintain strict separation between training and test datasets, regression coefficients for the covariates learned from the training sample were applied to the test sample to calculate effect size measures (Pearson's $\text{correlation}_{\text{cross-validated}}$). This procedure is described in detail in our previous publications (15,16).

We assessed the significance of all cross-validation-based correlations with non-parametric permutation tests. We randomly permuted the 5,821 subjects' outcome variable values 10,000 times and reran the PCR predictive modeling stream at each iteration, yielding a null distribution of correlation values. The procedure of Freedman and Lane (17) was used to account for covariates. In addition, exchangeability blocks were used to account for twin, family, and site structure and were entered into Permutation Analysis of Linear Models (PALM) (18) to produce permutation orderings.

#### 5. Latent Variable Modeling for Socioeconomic Resources

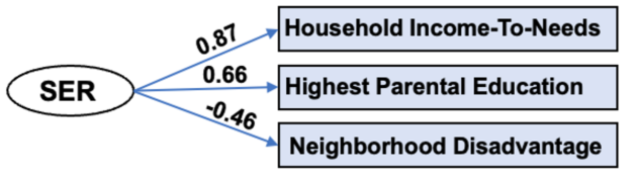


***Figure S3: Factor Model of Socioeconomic Resources.*** *Path estimates reflect standardized factor loadings.*

We constructed a latent variable for socioeconomic resources by applying factor analysis to household income-to-needs, highest parental education, and neighborhood disadvantage in 10,578 participants that had all three variables (see Figure S3). The latent variable accounted for 47% of the variance in the observed scores. Household Income-to-Needs combines information on household income, poverty lines, and family size. Household Income covered all sources of income for family members, including wages, benefits, child support payments, and others. It was assessed in bins as follows: 1 = <5,000, 2 = 5,000 - 11,999, 3 = 12,000 - 15,999, 4 = 16,000 - 24,999, 5 = 25,000 - 34,999, 6 = 35,000 - 49,999, 7 = 50,000 - 74,999, 8 = 75,000 - 99,999, 9 = 100,000 - 199,999, 10 = More than 200,000, and we assigned each subject the midpoint for their bin. Household size was calculated from the ABCD Parent Demographic Survey. The poverty line was calculated for a household of that size based on the 2021 US Poverty Guidelines ($8,340 + $4,540 per person in the household). Finally, Household Income-to-Needs was calculated as the ratio of the combined income midpoint over the poverty line. Highest Parental Education was the average educational achievement of parents or caregivers. Neighborhood disadvantage scores reflect an ABCD consortium-supplied variable (reshist_addr1_adi_wsum). Participant's primary home address was used to generate Area Deprivation Index (ADI) values (19), which were weighted based on results from Kind *et al.* (20) to create an aggregate measure. Higher scores on the factor indicate greater neighborhood disadvantage including higher percent of families living in poverty, increased unemployment, and lower levels of educational attainment at the neighborhood level.

#### 6. Analysis Using Both Positive and Negative Edges

Since most graph theory measures require unsigned edge weights, each subject's connectome resulted in two separate sets of graphs – one for the collection of positive edges and another for the negatively weighted edges (21,22). We focused on positive graphs exclusively because previous studies showed they contain the bulk of discriminative information about phenotypes. We also performed an additional analysis in which we calculated nodewise measures of within-module degree and participation coefficient for both positive and negative edges, resulting in 1,672 metrics in total. We used these metrics as predictive features in a new principal components regression analysis with SER as the outcome variable. We found the LOSO-CV out-of-sample correlation between predicted and actual SER scores was $r_{cv} = 0.142$, which is not meaningfully different than the correlation using positive edges alone ($r_{cv} = 0.162$).
